# The RAC1 activator Tiam1 regulates centriole duplication through controlling PLK4 levels

**DOI:** 10.1101/2020.05.20.106120

**Authors:** Andrew P. Porter, Gavin R. M. White, Erinn-Lee Ogg, Helen J. Whalley, Angeliki Malliri

## Abstract

Centriole duplication is tightly controlled to maintain correct centriole number through the cell cycle. A key component of this control is the regulated degradation of PLK4, the master regulator of centriole duplication. Here we show that the Rac1 guanine nucleotide exchange factor (GEF) Tiam1 localises to centrosomes during S-phase, where it is required for maintenance of normal centriole number. Depletion of Tiam1 leads to an increase in centrosomal PLK4, centriole overduplication and ultimately to lagging chromosomes at anaphase and aneuploidy. The effects of Tiam1 depletion can be rescued by re-expression of wild-type Tiam1 and catalytically inactive (GEF*) Tiam1, but not by Tiam1 mutants unable to bind to the F-box protein βTRCP, implying that Tiam1 regulates PLK4 levels through promoting βTRCP-mediated degradation.

## Results

Centrioles are barrel-shaped, microtubule-rich organelles fundamental to cell division, polarity and signalling. Centriole pairs recruit pericentriolar material (PCM) to form the centrosome, the major microtubule organising centre [1]. In G1, a typical human cell contains a single centrosome formed around two centrioles. At the G1/S transition, each of the mother centrioles generates a single daughter centriole (see schematic, Supplementary Figure 1a), which together recruit PCM to form one of the mitotic spindle poles [2].

**Figure 1.**
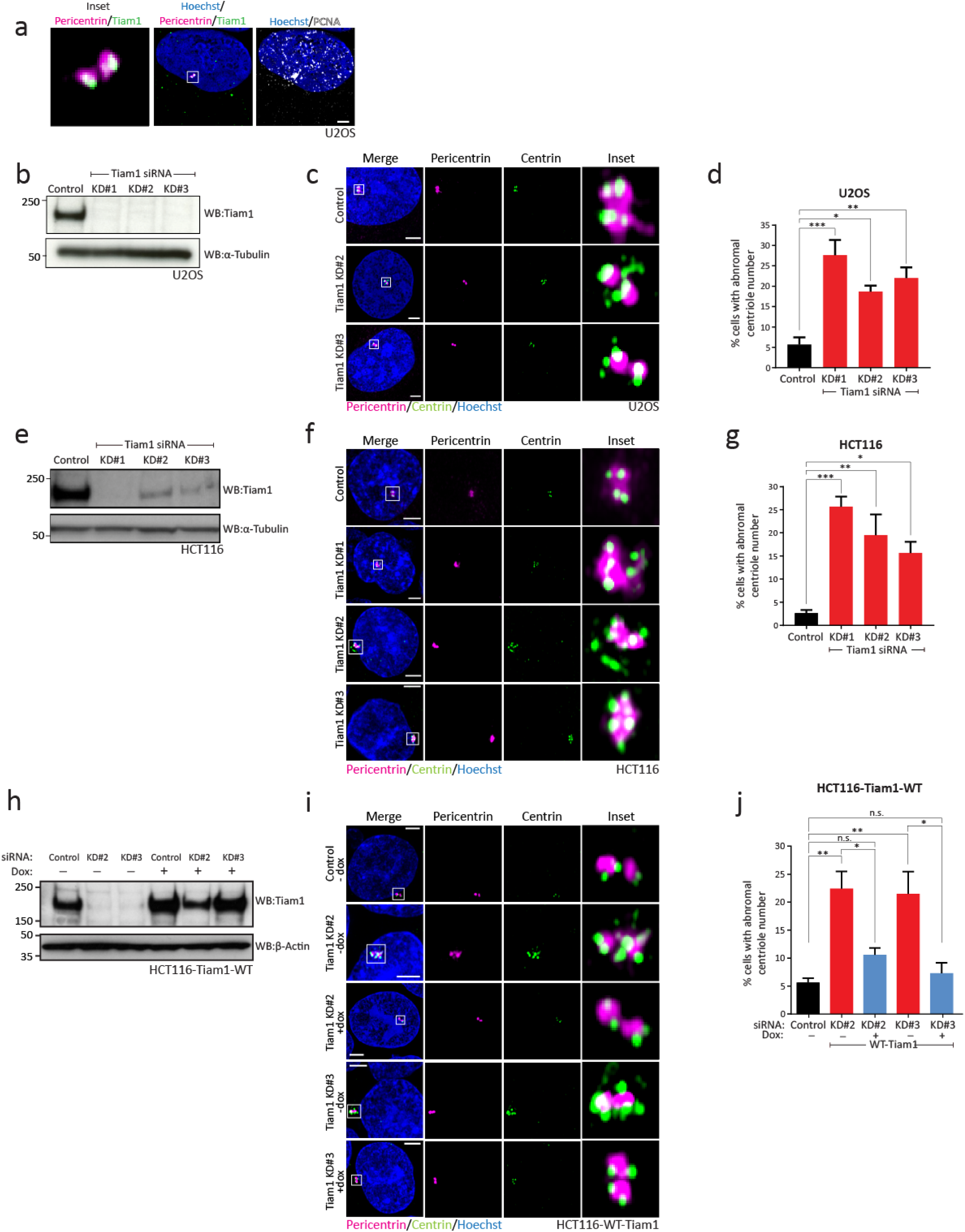
Tiam1 is required for maintenance of normal centriole number. a) Endogenous Tiam1 (green) localises to centrosomes (marked by Pericentrin, magenta) during S-phase (identified by punctate nuclear staining of PCNA, grey). Maximal z-projection of planes containing Pericentrin staining. b) Western blot showing depletion of endogenous Tiam1 in U2OS cells following transfection with three independent siRNAs. c) Maximal z-projections of confocal images showing normal centriole number in U2OS cells transfected with control siRNA, and excess centriole number in Tiam1-knockdown cells. Individual centrioles marked with Centrin (green), centrosomes marked with Pericentrin (magenta). d) Quantification of U2OS cells with abnormal centriole number from 3 independent experiments, quantified from images as in (c); more than 100 cells counted per condition per experiment (one-way ANOVA, corrected for multiple comparisons.) e) Western blot showing depletion of endogenous Tiam1 in HCT116 cells following transfection with three independent siRNAs. In (b) and (e) α-Tubulin was used as a loading control. f) Representative confocal microscopy images showing normal centriole number in HCT116 cells transfected with control siRNA, and excess centriole number in Tiam1-knockdown cells. Individual centrioles marked with Centrin (green), centrosomes marked with Pericentrin (magenta). g) Quantification of HCT116 cells with abnormal centriole number from 3 independent experiments (except Tiam1 KD#2, n=2), quantified from images as in (f); more than 100 cells counted per condition per experiment (one-way ANOVA, corrected for multiple comparisons.) h) Western blot from HCT116-Tiam1-WT cells showing depletion of endogenous Tiam1 following siRNA transfection with Tiam1 KD#2 and KD#3, and restoration of near-endogenous levels of Tiam1 following treatment with doxycycline (Dox) to induce expression of siRNA-resistant wild-type (WT) Tiam1. β-Actin was used as a loading control. i) Representative confocal images of control, Tiam1 knockdown and rescue cells showing abnormal centriole number following Tiam1 knockdown and normal centriole number in cells re-expressing wild-type Tiam1. j) Quantification of images as in (i) from 4 independent experiments; more than 50 cells counted per condition per experiment (one-way ANOVA, corrected for multiple comparisons; t-test for comparing +/-dox cells treated with the same siRNA.) For all quantification: *** p<0.001 ** p<0.01 * p<0.05 n.s. = not significant; error bars show S.E.M. All scale bars 3µm.

Centriole number is tightly controlled. Increased centriole or centrosome number leads to severe mitotic aberrations [2, 3]. During mitosis, cells with centrosome amplification form transient multipolar spindles. Division can proceed in such cells through clustering of extra centrosomes to pseudo-bipolar spindles. However, this gives rise to lagging chromosomes and ultimately aneuploidy [4]. Moreover, even in cells with two centrosomes, uneven centriole number at the mitotic poles can lead to lagging chromosomes and aneuploidy [5]. In turn, aneuploidy leads to transcriptional changes that can affect cell growth or cause proteotoxic stress. Aneuploidy is also a feature of chromosomal instability (CIN), a driver of cancer [3]. Indeed, centrosome amplification alone is sufficient for the initiation of tumourigenesis [6].

To ensure correct centriole number, cells tightly regulate centriole production [2, 7]. PLK4, a member of the polo-like kinase family, is the master regulator of centriole biogenesis [7]. During the G1-S transition, PLK4 phosphorylates its partner STIL, increasing recruitment of SAS6, a key structural component of the new procentriole (Supplementary Figure 1a) [7]. This process is restricted by PLK4 degradation, which is mediated by the SCF E3 ligase complex. PLK4 proteins homodimerise and trans-autophosphorylate a phospho-degron recognised by βTRCP, an F-box protein that targets SCF to its substrates [8-10]. Thus, a transient increase in PLK4 activity is tightly coupled to its destruction, efficiently terminating its activity and preventing extra rounds of centriole duplication [9, 10]. Excess PLK4 has been shown to lead to centriole overduplication across multiple species and cell types [8, 9, 11-13].

Tiam1, a guanine nucleotide exchange factor (GEF) for the small GTPase Rac1 has a wide variety of functions including roles in cell adhesion, polarity, migration, transcription and tumourigenesis [14-19]. We previously showed that Tiam1 localises to centrosomes in prophase and prometaphase, where it activates Rac1 signalling to antagonise centrosome separation [20]. Centrosomal Tiam1 is phosphorylated by CDK1 and this is required for phosphorylation and consequently activation of the Rac1 effector PAK1/2 [21]. We decided to investigate whether Tiam1 localises to centrosomes in other phases of the cell cycle. We used our previously validated staining protocol [20] and stained U2OS cells with antibodies against Tiam1, Pericentrin (as a centrosome marker) and PCNA which produces punctate nuclear staining during S-phase [22]. We observed localisation of Tiam1 at centrosomes during S-phase (Figure 1a), as well as throughout mitosis as we previously published [20] (data not shown).

We hypothesized that Tiam1 plays an additional role in centrosome biology in S-phase, when centriole duplication occurs [7]. To test this, we depleted Tiam1 from U2OS cells using RNAi (Figure 1b). Control and knockdown cells were stained with antibodies against Pericentrin and Centrin (to mark individual centrioles) (Figure 1c). Cells depleted of Tiam1 displayed an increase in centriole number, with many cells containing centrosomes with additional Centrin puncta (i.e. greater than 2 centrioles per centrosome) (Figure 1c). Using three independent siRNA sequences targeting Tiam1, we saw a significant increase in the number of cells with centriole overduplication (Figure 1d).

To determine whether this effect could be observed in other cell lines, we used siRNA to deplete Tiam1 in HCT116 colon cancer cells, which, like U2OS cells, have a normal centriole number [5, 23]. Tiam1 depletion (Figure 1e) again resulted in excess centrioles compared with control siRNA (Figure 1f and 1g). We also observed similar effects in the human breast cancer cell line MCF7 (Supplementary Figure 1b-d). Together, this suggests a conserved role for Tiam1 in centriole duplication across multiple cell types. To determine whether these extra Centrin puncta represented true centrioles or rather centriolar satellites (which localise around the centrosome and stain with Centrin antibody [24]), we placed control and Tiam1 knockdown cells on ice for 30 minutes to depolymerise microtubules (Supplementary Figure 1e). While this causes dispersion of centriolar satellites, true centrioles are unaffected [25, 26]. We observed a significant increase in Tiam1 knockdown cells with excess centrioles following cold treatment (Supplementary Figure 1e and 1f), indicating that the extra Centrin puncta are indeed centrioles. As a further confirmation, we also looked specifically at mitotic U2OS cells and observed centriole overduplication at the mitotic poles following Tiam1 depletion, consistent with these aberrant centrioles forming functional centrosomes (Supplementary Figure 1g-1i).

To demonstrate that the effects of Tiam1 depletion are on-target effects of siRNA knockdown, we developed a system to restore wild-type Tiam1 levels using siRNA-resistant Tiam1. We constructed HCT116 cells expressing mouse WT-Tiam1 [intrinsically resistant to two siRNAs targeting the human Tiam1 sequence (Tiam1 KD#2 and Tiam2 KD#3)] under the control of a doxycycline-inducible promoter, as in previous publications [17, 21]. We could detect the exogenous WT-Tiam1 localising to centrosomes by immunofluorescence following doxycycline treatment (Supplementary Figure 1j). As before, siRNA treatment effectively depleted endogenous Tiam1; however, addition of doxycycline led to near-endogenous expression of WT-Tiam1 protein (Figure 1h). Tiam1 depletion again led to an increase in centriole number, which was effectively rescued by doxycycline-induced WT-Tiam1 (representative images in Figure 1i, quantified in 1j). This demonstrates that Tiam1 is required for normal centriole number.

To determine the mechanism by which Tiam1 maintains normal centriole number, we first used flow cytometry to investigate whether Tiam1 knockdown affected cell cycle progression, as increased S-phase duration can lead to centriole overduplication [27], while failure to undergo cytokinesis results in centrosome amplification [28]. However, we saw no difference in cell cycle progression nor evidence of polyploidy comparing control and Tiam1 knockdown cells (Supplementary Figure 2a-2c). Having excluded a cell cycle defect, we decided to next examine whether Tiam1 knockdown affected PLK4 levels at centrosomes, PLK4 being the key driver of centriole duplication [8, 9, 11-13]. We first confirmed that our antibody detected centrosomal PLK4 by staining U2OS cells treated with either a control siRNA or several PLK4 siRNAs (Supplementary Figure 2d). We calculated PLK4 centrosomal intensity (Supplementary Figure 2e) and saw a decrease in centrosomal PLK4 in all siRNA-treated cells compared with control cells (Supplementary Figure 2f).

Next we stained control or Tiam1-depleted U2OS cells for PLK4 and Pericentrin. We saw a significant increase in centrosomal PLK4 in cells treated with Tiam1 siRNA compared with control siRNA-treated cells (Figure 2a, quantified in 2b). The equivalent experiment with HCT116 cells also demonstrated an increase in centrosomal PLK4 staining after Tiam1 depletion (Figure 2c, quantified in 2d).

**Figure 2.**
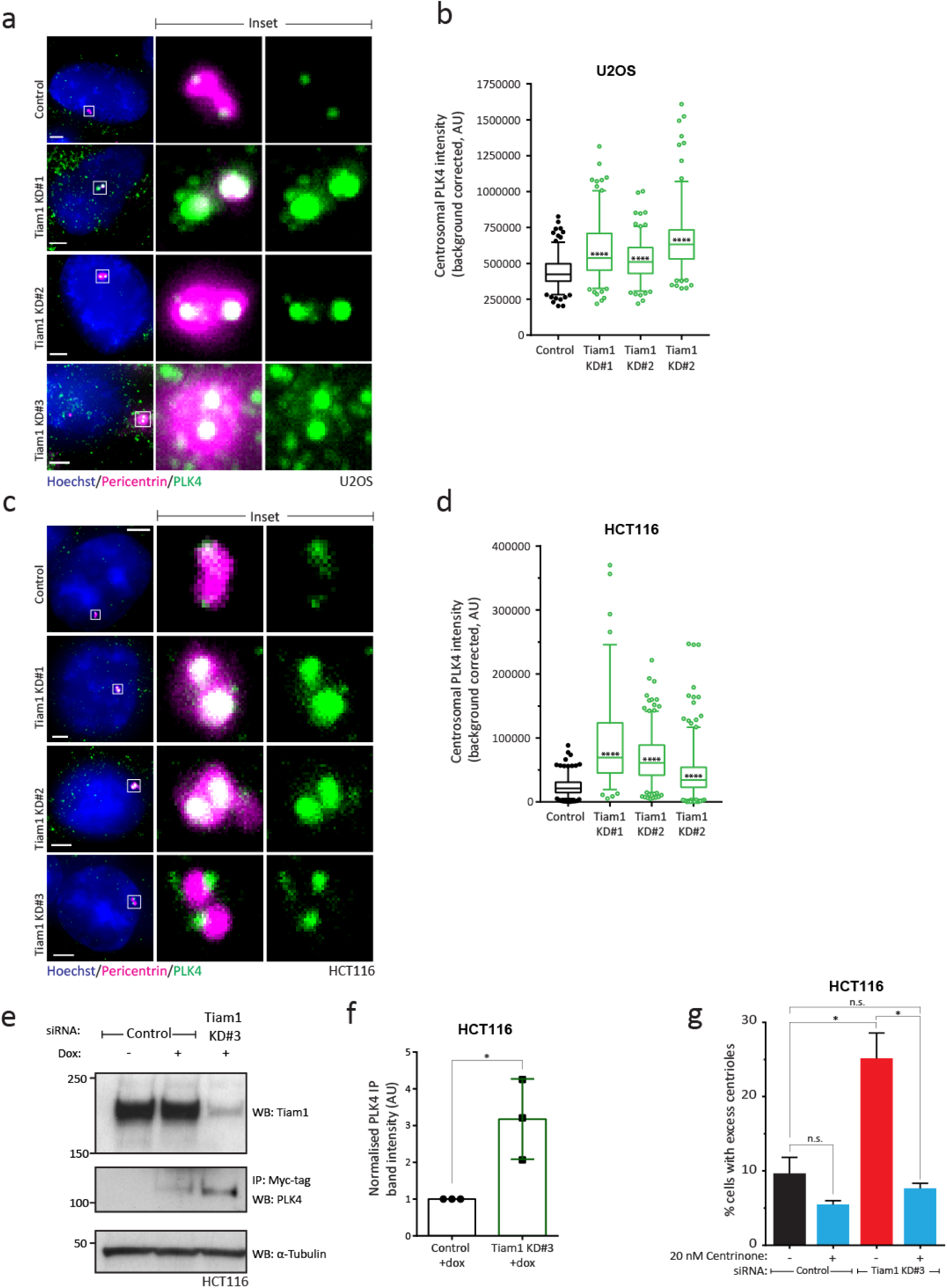
Tiam1 knockdown leads to an increase in centrosomal PLK4. a) Maximal z-projections of U2OS cells stained for Pericentrin (magenta) as a marker of centrosomes, and endogenous PLK4 (green), showing an increase in centrosomal PLK4 staining intensity in Tiam1-knockdown cells. b) Quantification of PLK4 intensity at centrosomes in U2OS cells from three independent experiments from images as in (a) [total number of cells: n=186 (control), n=175 (KD#1), n=165 (KD#2), n=170 (KD#3)]. Box shows 25th to 75th percentiles, whiskers show 5th to 95th percentiles; median is marked with a line. One-way ANOVA, corrected for multiple comparisons. c) Maximal z-projections of HCT116 cells stained for Pericentrin (magenta) as a marker of centrosomes, and endogenous PLK4 (green), showing an increase in centrosomal PLK4 staining in Tiam1-knockdown cells. d) Quantification of PLK4 intensity at centrosomes in HCT116 cells from two independent experiments from images as in (c) [total number of cells: n=237 (control), n=97 (KD#1), n=251 (KD#2), n=296 (KD#3)]. Box shows 25th to 75th percentiles, whiskers show 5th to 95th percentiles; median is marked with a line. One-way ANOVA, corrected for multiple comparisons. e) Representative western blot showing an increase in expression of exogenous myc-tagged PLK4 (as determined by immunoprecipitation of myc-tagged PLK4) in U2OS cells following Tiam1 knockdown, compared with control transfected cells. Dox=doxycycline, added to induce expression of exogenous myc-tagged PLK4. α-Tubulin was used as a loading control. f) Quantification of increased PLK4 expression (as determined by immunoprecipitation of myc-tagged PLK4) from control and Tiam1 knockdown cells (n=3 independent experiments, bars show mean +/-SEM, dots represent each individual experiment). PLK4 pull-down levels normalised to Tubulin input. g) Quantification of cells with abnormal centriole number in U2OS cells transfected with either control siRNA or Tiam1 KD#3 siRNA, and treated with either vehicle or Centrinone (20 nM). For all quantification: **** p<0.0001 * p<0.05. Scale bars are 3µm.

To corroborate these immunofluorescence data, we biochemically measured PLK4 levels following Tiam1 knockdown. We were unable to detect endogenous PLK4 by western blot (similar to reports from other researchers, as the cellular levels of PLK4 are very low [10]), and therefore over-expressed myc-tagged PLK4 in U2OS cells using a doxycycline-inducible system. This induced centriole overduplication (Supplementary Figure 2g). We used anti-Myc beads to immunoprecipitate exogenous PLK4, and saw that its levels increased following Tiam1 knockdown (Figure 2e, quantified in 2f). Together, these experiments indicate that Tiam1 limits PLK4 levels at the centrosome, thereby constraining centriole duplication.

Given that Tiam1-depleted cells have elevated PLK4 levels and that high PLK4 drives centriole overduplication, we hypothesised that low-level PLK4 inhibition in Tiam1 depleted cells would restore their correct centriole number. We therefore treated HCT116 cells with the specific PLK4 inhibitor Centrinone [29] at 20 nM, a concentration which had minimal effects on centriole number in control siRNA treated cells (Figure 2g). Tiam1 knockdown in the absence of the inhibitor led to an increase in centriole number (Figure 2g), whereas knockdown cells treated with Centrinone did not display an increase in centriole number compared to control-siRNA treated cells (Figure 2g). These findings show that the production of extra centrioles in Tiam1-depleted cells requires increased PLK4 activity.

We next investigated the mechanism by which Tiam1 regulates PLK4 levels. Since Tiam1 is best known for its ability to activate Rac1, we addressed whether the catalytic (GEF) activity of Tiam1 was required for correctly regulating centriole numbers. We therefore expressed a doxycycline-inducible siRNA-resistant ‘GEF-dead’ mutant (GEF*) Tiam1 (Figure 3a) in HCT116 cells treated with either control siRNA or siRNA against endogenous Tiam1 (Figure 3b). GEF* Tiam1 localised to the centrosomes similarly to wild-type Tiam1 (Supplementary Figure 3a) and its expression restored normal centriole number in cells depleted of endogenous Tiam1 (Figure 3c, quantified in 3d). This indicates that Tiam1-dependent Rac1 activation is not required for normal centriole duplication nor its localisation to the centrosome. This also suggests that this pathway is distinct from the one controlling centrosome separation during prophase and prometaphase, which does rely on Tiam1-mediated Rac1 activation [20, 21]. To further investigate the relationship between these two centrosomal Tiam1 functions, we performed rescue experiments with Tiam1-1466A, a CDK1-phosphorylation mutant version of Tiam1 (Supplementary Figure 3b) that is unable to rescue centrosome separation in prophase despite correct localisation to centrosomes [21].

**Figure 3.**
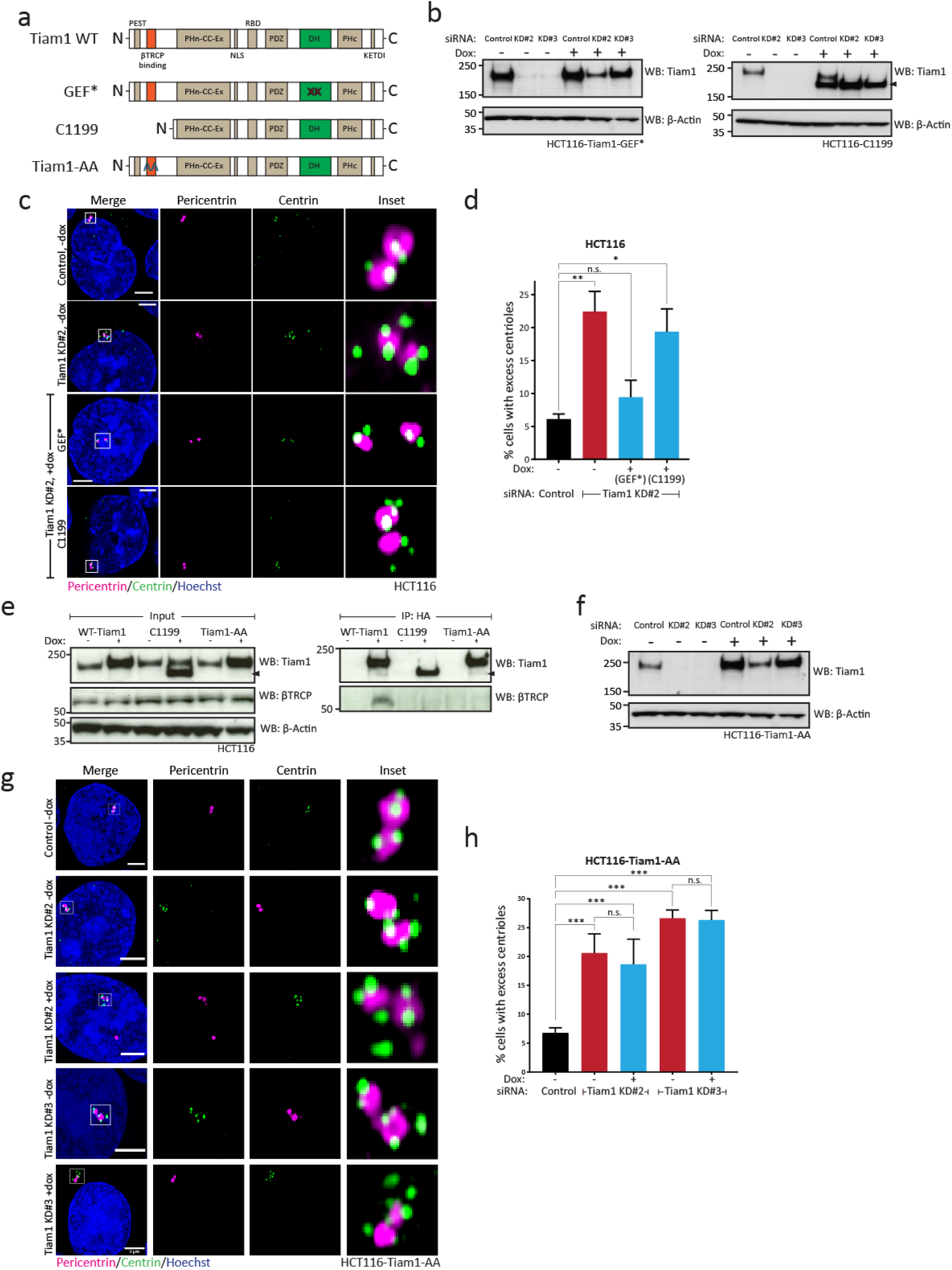
Tiam1 requires βTRCP binding to maintain normal centriole number. a) Schematic showing the domain structure of wild-type (WT) Tiam1, and the mutant versions. b) Western blots from HCT116-Tiam1-GEF* and HCT116-C1199 cells showing depletion of endogenous Tiam1 by two independent siRNAs and, following treatment with doxycycline (Dox), restoration of near-endogenous levels of either a GEF-dead (GEF*) Tiam1 or an N-terminally truncated mutant (C1199, marked with an arrowhead), both of which are resistant to siRNA treatment. c) Maximal z-projection confocal images of HCT116 control, Tiam1 knockdown and Tiam1-rescue cells. Cells exhibit abnormal centriole number following Tiam1 knockdown; this is rescued by expression of RNAi-resistant GEF* Tiam1, but not by expression of RNAi-resistant C1199. d) Quantification of HCT116 cells with excess centrioles from rescue experiments as in (c), from 3 independent experiments; more than 50 cells counted per condition per experiment (one-way ANOVA, corrected for multiple comparisons). e) Western blot from HCT116 cells showing immunoprecipitation of wild-type Tiam1, C1199 (marked with an arrowhead) and Tiam1-AA (all constructs HA-tagged); only wild-type Tiam1 is able to co-immunoprecipitate endogenous βTRCP. f) Western blot from HCT116-Tiam1-AA cells showing depletion of endogenous Tiam1 by two independent siRNAs and restoration of near-endogenous levels of Tiam1 following treatment with doxycycline (Dox) to express the RNAi-resistant, non-βTRCP binding mutant Tiam1-AA. g) Maximal z-projection confocal images of control and Tiam1-knockdown HCT116 cells, showing excess centriole number following Tiam1 knockdown, which is not rescued by doxycycline-induced expression of siRNA-resistant Tiam1-AA. h) Quantification of cells with excess centrioles from rescue experiments as in (g) from 4 independent experiments; more than 50 cells counted per condition per experiment (one-way ANOVA, corrected for multiple comparisons; t-test for comparing +/-dox cells treated with the same siRNA). For all quantification: *** p<0.001 ** p<0.01 * p<0.05; error bars show S.E.M. Scale bars show 3µm. β-Actin was used as a loading control in all blots.

As with the GEF* mutant, expression of this mutant Tiam1 was able to restore normal centriole number after Tiam1 knockdown (Supplementary Figure 3c and 3d), demonstrating that this role of Tiam1 is distinct from that of Tiam1 in prophase.

As the function of Tiam1 in centriole duplication appeared to be independent of Rac1, some other domain of Tiam1 must play a role in regulating centriole duplication. Indeed, Tiam1 is a large multi-domain protein, which also acts as a molecular scaffold [30]. We therefore performed our rescue experiment again with a well-characterised, N-terminally truncated mutant of Tiam1, C1199 (Figure 3a). Interestingly, doxycycline-induced expression of siRNA-resistant C1199 (Figure 3b) was unable to rescue centriole number in HCT116 cells (Figure 3c, quantified in 3d). This suggests that a function performed by the N-terminus of Tiam1 is required for normal centriole duplication. Significantly, this region of Tiam1 contains a phospho-degron required for binding of βTRCP [31], a component of the E3-ligase complex also targeting centrosomal PLK4 for degradation during S-phase [8, 32].

We reasoned that the interaction between Tiam1 and βTRCP may be required for the ability of Tiam1 to regulate PLK4 levels at the centrosome, and that the C1199 mutant may be unable to substitute for endogenous Tiam1 because of its lack of βTRCP binding. To test this, we produced a mutant version of full-length Tiam1 (referred to as Tiam1-AA; Figure 3a) containing two point mutations in the βTRCP phospho-degron [31]. We confirmed that both this mutant and C1199 were unable to bind to βTRCP by co-immunoprecipitation (Figure 3e). We also confirmed that Tiam1-AA was able to correctly localise to the centrosome in the same way as wild-type Tiam1 (Supplementary Figure 3a), indicating that βTRCP binding was not required for centrosomal localisation of Tiam1. We then performed Tiam1 knockdown and rescue experiments, inducibly expressing the siRNA-resistant Tiam1-AA mutant to near-endogenous levels in cells depleted of Tiam1 (Figure 3f). In both uninduced (-dox) and induced (+dox) cells there was a significant and indistinguishable increase in cells with excess centrioles upon endogenous Tiam1 depletion compared with control-siRNA treated cells (Figure 3g, 3h), showing that expression of Tiam1-AA is unable to compensate for depletion of endogenous Tiam1. Therefore, we conclude that regulation of normal centriole number requires the interaction between Tiam1 and βTRCP.

Centriole overduplication can lead to chromosome mis-segregation, aneuploidy and chromosomal instability (CIN) [33], ultimately leading to tumourigenesis [6, 12, 34]. Chromosome mis-segregation can occur either through an imbalance of centriole numbers at the two poles of the mitotic spindle [5] or via the formation of transient multipolar intermediates arising from centrosome amplification [4]. Depletion of Tiam1, apart from triggering centriole overduplication, also led to low levels of centrosome amplification (Supplementary Figure 4a and 4b). Interestingly, Tiam1 depletion, while largely suppressing tumour formation [14, 16], promotes malignant progression [14, 16] indicating a dual oncogene/tumour suppressor role for Tiam1. As CIN is considered to drive the acquisition of malignant hallmarks [35, 36], we investigated whether knockdown of Tiam1 could lead to an increase in lagging chromosomes at anaphase, a widely used readout of chromosome mis-segregation. We combined Hoechst staining with a centromere marker (CREST) to distinguish true lagging chromosomes from acentromeric chromosome fragments and chromosome bridges (Figure 4a). In the chromosomally-stable HCT116 cell line, we observed a low level of lagging chromosomes at anaphase and early telophase in control-siRNA treated cells (Figure 4a, quantified in 4b). Following Tiam1 knockdown, the number of cells with lagging chromosomes significantly increased (Figure 4a, 4b). There was no change in the number of chromosome bridges or acentric chromosomes (data not shown), suggesting that this phenotype affects the segregation of whole chromosomes specifically. We saw similar increases in lagging chromosomes in U2OS cells depleted for Tiam1 (Supplementary Figure 4c, quantified in 4d), indicating conservation of the role of Tiam1 in chromosome segregation.

**Figure 4.**
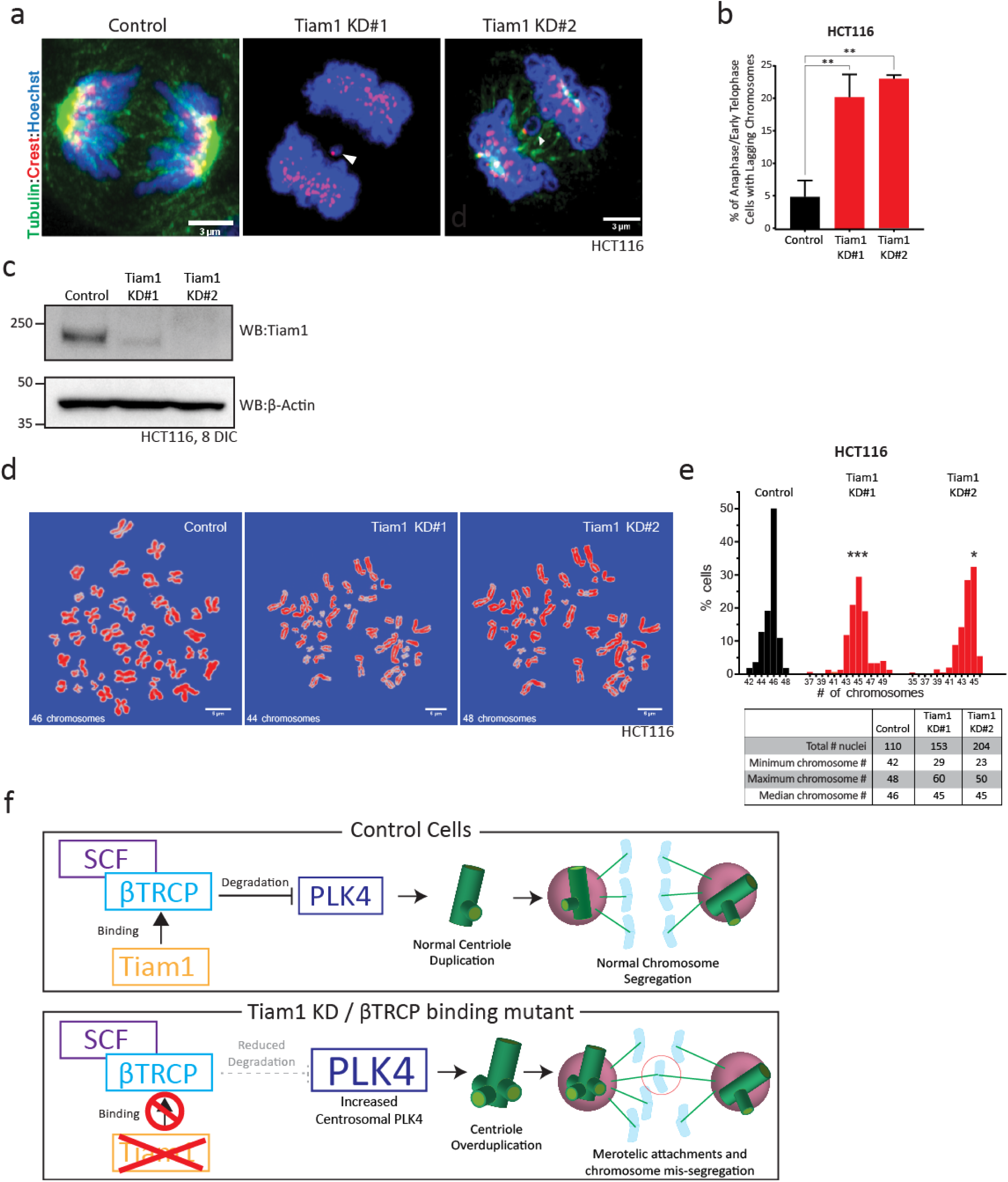
Depletion of Tiam1 leads to lagging chromosomes at anaphase and early telophase and aneuploidy. a) Maximal z-projection confocal images of anaphase HCT116 cells; chromosomes stained with Hoechst and marked with CREST as a centromere marker, showing lagging chromosomes following Tiam1 knockdown with two independent siRNAs. Scale bars are 3µm. b) Quantification of percentage of HCT116 cells in anaphase or early telophase with lagging chromosomes following transfection with control or Tiam1 knockdown siRNAs, as in (a). Data quantified from three independent experiments, at least 50 cells quantified per condition per experiment (one-way ANOVA, corrected for multiple comparisons). c) Western blot showing Tiam1 levels in HCT116 cells following 8 days growth in culture with repeated siRNA transfection. β-Actin was used as a loading control. d) Representative images of metaphase spreads from HCT116 cells treated with either control or Tiam1 siRNAs. Scale bars are 5µm. e) Histogram of chromosome number from control and Tiam1 knockdown HCT116 cells; data collected from three independent experiments. Table shows additional statistics. Kruskal-Wallis test adjusted for multiple comparisons. * p<0.05 *** p<0.001. f) Model of how the Tiam1-βTRCP interaction affects PLK4 protein levels, centriole duplication and chromosome segregation.

Given this increase in lagging chromosomes, and the previously reported chromosome alignment defects arising from Tiam1 knockdown [20], we tested whether these would translate into an increase in aneuploidy given longer periods of Tiam1 depletion. We grew HCT116 cells for 8 days with two rounds of siRNA transfection (Figure 4c), and prepared metaphase spreads. Chromosome number was determined by manual counting of Hoechst-stained chromosomes (representative images in Figure 4d). While the range of chromosome number found in control cells was relatively narrow (from 42 to 48 chromosomes), we found a much wider distribution of chromosome number in cells treated with Tiam1 siRNA (from 29 to 60 for Tiam1 KD#1, and from 23 to 50 for Tiam1 KD#2), indicating a significant increase in aneuploidy following Tiam1 depletion (Figure 4e). Our data indicate that Tiam1 contributes to the control of chromosome segregation through regulating centriole duplication.

## Discussion

In this study, we identify Tiam1 as a new regulator of PLK4, the master regulator of centriole duplication. PLK4 regulation involves a fine balance between a brief period of activity – sufficient to initiate procentriole formation – and ubiquitin-mediated degradation to prevent re-duplication. Too much activity leads to overduplication [1, 8, 9], too little to centriole loss [29], but the full details of this precise spatio-temporal regulation remains to be determined [7]. Defects in PLK4 levels can ultimately lead to defects in chromosome segregation [4, 6] and other pro-oncogenic effects such as increased invasion [28, 37].

Our data indicate that Tiam1 acts as a modulator of PLK4 protein levels, with depletion of Tiam1 leading to an increase in centrosomes containing more than 2 centrioles, and a small increase in cells with centrosome amplification (see model in Figure 4f). While we have yet to establish how precisely Tiam1 modulates PLK4 levels through βTRCP, in a previous publication, we showed that the interaction between βTRCP and Tiam1 was necessary for the degradation of TAZ, another βTRCP target protein [19]. Given our current findings, this raises the possibility that Tiam1 is able to act as a scaffold for βTRCP more generally, directing it to its targets, or enhancing its interaction with target proteins. As Tiam1 is itself a target of βTRCP-mediated degradation, this could allow for precise temporal control of βTRCP-target degradation, as once Tiam1 itself is degraded the targeting effect would be removed.

The regulation of centriole duplication by Tiam1 appears distinct from its role in driving centrosome separation during prophase [20], as, unlike the latter, the former does not depend on activation of Rac1, nor does it require phosphorylation of Tiam1 at S1466. It is interesting to speculate whether these two centrosomal functions of Tiam1 are temporally segregated, and if so how. Perhaps distinct complexes of Tiam1 exist at centrosomes (including either βTRCP and PLK4 or Rac1 and PAK1/2) dictated by mutually exclusive intermolecular interactions. Alternatively, post-translational modification of Tiam1, such as by CDK1 phosphorylation [21], might control a temporal switch between interacting partners.

With respect to the impact of Tiam1 depletion on chromosomal stability, we show here that loss of Tiam1 leads to an increase in lagging chromosomes at anaphase and to aneuploidy. The latter is likely due to a combination of the effects of centriole overduplication and centrosome amplification demonstrated in this study, and chromosome congression defects that arise from Tiam1 depletion, as we previously published [20, 21]. Together these indicate a pathway by which Tiam1 depletion could enhance tumour progression. While targeting the Tiam1-Rac pathway therapeutically remains a subject of ongoing investigation, due to the dramatic reduction in tumour formation following loss of Tiam1 in animal models [14, 16], this work highlights a need for more detailed understanding of Tiam1 signalling to separate the pro- and anti-tumourigenic properties of Tiam1.

## Materials and Methods

### Cell Culture

All cell lines were cultured at 37°C in a humidified incubator (5% CO2 atmosphere). U2OS and HCT116 cells were cultured in DMEM High Glucose (Gibco) supplemented with 10% FBS (Gibco). MCF7 cells were cultured in DMEM High Glucose (Sigma) supplemented with 10% FBS, 1% L-Glutamine (Gibco). Cell lines were routinely tested to exclude Mycoplasma contamination and for cell line authentication (via in-house facilities).

### Generation of cell lines

Plasmids were introduced into cells either by transfection using TransIT-LT1 (Mirus) according to the manufacturer’s instructions or by retroviral transduction as previously described [20]. For inducible overexpression, HCT116 were retrovirally transduced with pRetro-Tet-ON followed by selection with G418 (1mg/ml, Sigma-Aldrich). pRetro-XT-based constructs were then retrovirally transduced and cells selected with puromycin (2 μg/ml, Sigma-Aldrich).

### Plasmids

The doxycycline-inducible PLK4 plasmid was generated by PCR of the wild-type PLK4 cDNA from Addgene plasmid #41165 by PfuUltra II Fusion HS DNA polymerase (Agilent #600670.) pcDNA Plk4(Sak) wt (Nigg HR9) was a gift from Erich Nigg[38]. The sequence was confirmed by PCR, and digested with NotI and MluI enzymes (restriction sites introduced via the PCR primer sequences) for introduction into similarly-digested pRetro-X-Tight (Takara Bio). The final insertion was confirmed by sequencing.

The doxycycline-inducible Tiam1-AA (non-βTRCP-binding) mutant was generated by Quikchange II (Agilent) site-directed mutagenesis of WT-Tiam1 mouse cDNA at S329 and S334, and successful mutagenesis confirmed by PCR. A portion of the cDNA containing the mutated region was digested with HpaI and NotI, and inserted into the existing pRetro-X-Tight-Tiam1-WT plasmid. Insertion was confirmed by PCR.

### siRNA transfection

For transfection of cells for immunofluorescence staining, cells were plated at a density of 2×10^5^ onto glass coverslips (6 well dish), and reverse transfected using RNAiMax (Invitrogen) following the manufacturer’s instructions. Cells were grown for 72 hours before being fixed in MeOH. For Tiam1, four siRNA sequences were used - Tiam1 KD#1 5’-GAGGTTGCAGATCTGAGCA-3’; Tiam1 KD#2 5’-GAGGUUGCAGAUCUGAGCA-3’; Tiam1 KD#3 5’-AGAGCGCACCUACGUGAAA-3’; Tiam1 KD#4 5’GGTTCTGTCTGCCCAATAA3’ - and synthesized by Eurofins MWG. In all of the reported assays, a negative control siRNA was used typically siLuc (control) 5′-CGUACGCGGAAUACUUCGA-3′, or Dharmacon siGENOME Non-Targeting siRNA #4. 4 siRNA sequences were used to deplete PLK4 to test the specificity of the anti-PLK4 antibody. These were: PLK4 siRNA#1: 5′-GAAAUGAACAGGUAUCUAA-3′; PLK4 siRNA#2: 5′-GAAACAUCCUUCUAUCUUG-3′; PLK4 siRNA#3: 5′-GUGGAAGACUCAAUUGAUA-3′; and PLK4 siRNA#4: 5′-GGACCUUAUUCACCAGUUA-3′ (sequences derived from [39]).

### Centrinone Treatment

Following siRNA transfection and plating, cells were treated with 20nM Centrinone (Tocris Bioscience) or vehicle control for 3 days before fixation.

### Depolymerisation of the cytoplasmic microtubule network

Tiam1 was transiently depleted in U2OS cells by reverse transfection of siRNA on glass coverslips. 72 hours post transfection cells were transferred to pre-cooled DMEM (4°C) and incubated on ice for 30 minutes to depolymerise cytoplasmic microtubules.

Cells were fixed in ice-cold methanol immediately following 30 minute incubation on ice. Centrioles were visualised by IF using Centrin and centrosomes using Pericentrin. Diffuse alpha-Tubulin staining around the centrioles revealed depolymerisation of the microtubule network. Centriole number was quantified using the Deltavision Core microscope.

### Antibodies

Antibodies against the following were used for western blotting (WB), immunofluorescence (IF) and immunoprecipitation (IP):

- α-tubulin (DM1A; Sigma-Aldrich, T6199 mouse; 1:2500 IF, methanol; 1:5000 WB)
- β-actin (Sigma-Aldrich, clone AC-15 mouse; 1:10,000 WB)
- Centrin (Millipore, (20H5) Mouse;1:5000 IF, methanol)
- CREST (Europa Bioproducts, FZ90C-CS1058, Human, 1:2000, methanol)
- HA tag (Roche Diagnostics, 3F10 rat; 1:200 IF, formaldehyde; 1:1000 WB)
- PCNA (Abcam, ab18197, rabbit, 1:2000 IF, methanol)
- Pericentrin (Covance, PRB-432C; rabbit, 1:2000 IF, methanol)
- PLK4 (Sigma-Aldrich, clone 6H5 mouse, 1:1000 IF, methanol; 1:500 WB)
- Tiam1 (Bethyl Laboratories, rabbit, A300-099A, 1:1000 WB)
- Tiam1 (R&D Systems, AF5038, sheep 1:200, methanol)

Secondary antibodies and stains used were: IgG peroxidase-conjugated anti-mouse IgG from donkey (GE Healthcare, NA931), anti-rabbit IgG from donkey (GE Healthcare, NA934); Alexa Fluor 488 chicken anti-mouse IgG (H+L) (Molecular Probes, A21200), Alexa Fluor 488 donkey anti-rat IgG (H+L) (Molecular Probes, A21208), Alexa Fluor 568 donkey anti-mouse IgG (H+L) (Molecular Probes, A10037), Alexa Fluor 647 chicken anti-rabbit IgG (H+L) (Molecular Probes, A21208) (1:500 IF); Alexa Fluor 647 goat anti-human IgG (H+L) (Molecular Probes, A21445) (1:500 IF); Hoechst 3342 (Life Technologies, H3570).

### Protein analysis

Cells were lysed in an appropriate volume of IP lysis buffer [50 mM Tris-HCl pH 7.5, 150 mM NaCl, 1% Triton-x-100 (v/v), 10% glycerol (v/v), 2 mM EDTA, 25 mM NaF and 2 mM NaH2PO4 containing 1% protease inhibitor cocktail (P8340, Sigma-Aldrich) and 1% phosphatase inhibitor cocktails 2 and 3 (P5726 and P0044, Sigma-Aldrich) added fresh] or RIPA buffer [25 mM Tris pH 7.5, 150 mM NaCl, 0.1% SDS (v/v), 0.5% sodium deoxycholate (v/v), 1% Triton-x-100 containing 1 EDTA-free protease inhibitor tablet (Roche) and 1% phosphatase inhibitor cocktails 1 and 2 (Sigma-Aldrich) added fresh] for 10 min on ice and proteins were resolved by SDS-PAGE for western blotting.

For immunoprecipitation of myc-tagged PLK4, lysates were incubated with 50 μL of Myc-tag beads (A7470, Sigma-Aldrich, blocked with 5% BSA for one hour at room temperature), for 2 hours at 4 °C with rotation. Beads were subsequently washed with lysis buffer, and eluted with 2× SDS-PAGE sample buffer (Nupage, Invitrogen).

### Immunofluorescence

For immunofluorescence, cells were grown on coverslips and fixed with 100% ice-cold methanol for 5 min at –20 °C. Cells were washed and then blocked in 1% BSA in PBS (v/v) for 1 hour, before successive incubation with primary antibodies (overnight at 4oC) and then secondary antibodies (1 hour at room temperature). Coverslips were mounted onto glass slides using Fluromount-G (Southern Biotech) along with Hoechst 33342 (1:5000) for nuclear staining, or droplet of ProLong® Gold anti-fade reagent containing the DNA stain DAPI. Staining of endogenous Tiam1 was performed with the same protocol as in [21] which was shown to specifically detect centrosomal Tiam1.

### Metaphase spreads

HCT116 cells were grown for 8 days, with an initial siRNA transfection followed by two subsequent trypsinisation, replating and transfections (total of three rounds of siRNA transfection). Cells were treated with 150nM Nocodazole (M1402, Sigma-Aldrich) for four hours to arrest cells in mitosis. Mitotic cells were collected by a mitotic shake off and harvested, followed by resuspension in 3ml pre-warmed hypotonic buffer (40% RPMI (R8758, Sigma-Aldrich), 60% ddH_2_O) for 20 minutes. Cells were fixed by repeated addition of Carnoy’s fixative (3:1 v/v solution of Methanol:Glacial Acetic Acid), centrifugation and aspiration before final collection in 100% glacial acetic acid. Cells were dropped onto precooled (4°C) wet slides from a height of 40-50cm, left to dry and stained with Hoechst. Images were taken using the Zeiss AiryScan confocal and chromosome number determined by manual counting using ImageJ.

### Microscopy

Centrosome and centriole images were acquired using a Zeiss Observer equipped with a Zeiss LSM 880 scan head with the AiryScan detector, with Argon laser 458, 488, 514nm (Lasos, Jena, Germany), Diode 405-30 (Lasos), DPSS 561-10 (Lasos) and HeNe 633nm (Lasos) were utilised for illumination and a Plan-Apochromat 40×/1.4 Oil (Zeiss) objective lens. Centriole, lagging chromosome and metaphase spread images were acquired utilising the ‘Fast’ AiryScan mode. All equipment control, acquisition and processing of AiryScan images was performed in Zen Black (Zeiss). All images were processed in ImageJ. Images in the same figure panel stained with the same antibody are all set to the same minimum and maximum brightness for comparison of localisation and intensity.

PLK4 images were captured using a Zeiss Axiovert 200 M microscope (Solent Scientific). The system uses an Andor iXon 888 camera and a 300 W Xenon light source is used for fluorescence illumination with a variety of ET-Sedat filters (406, 488, 568, 647 nm). The system utilises the Metamorph software to capture and process images. Images were taken using the 100 × oil lens.

Immunofluorescence images of centrosomes and centrioles in MCF7 cells and mitotic U2OS cells, as well as in cold-treated U2OS cells, were captured using the Deltavision Core system (based on an Olympus IX71 microscope; fluorescence is achieved using a 300 W Xenon light source with a variety of Sedat filter sets (406, 488, 568, 647 nm) and the attached Roper Cascade 512B camera; images were taken using 100 × / 60 × oil lens.) The Deltavision core system utilises softWorx to capture and process images.

### Centriole counts

Widefield or confocal z-stacks of cells stained for Centrin and Pericentrin were used to determine centriole number. Planes were taken at the minimum distance recommended by the system (typically either 200nm or 140nm, depending on the system). Centriole counts were conducted manually using ImageJ, with any cell containing at least one centrosome (marked with Pericentrin) which was associated with three or more puncta of Centrin counted as having excess centriole number. Also included were cells with more than two distinct centrosomes, as these were considered to have arisen from earlier centriole overduplication events (although these typically accounted for only 2-4% of abnormal cells).

### Quantification of centrosomal PLK4 intensity

Maximal projections of widefield microscopy image stacks were generated using ImageJ. Two square regions of interest were drawn of fixed size: the first encompassing the centrosome area as marked with Pericentrin (35px^2^) (a) and a second larger region of interested, centred on this first box (55px^2^) (b). The formula a - ((b-a)(55^2^/35^2^)) was used to calculate the intensity of the PLK4 signal at the centrosome, adjusted for background intensity surrounding the centrosome.

### Statistical analysis

Appropriate statistical tests were chosen to minimise type I error associated with significance values. Statistical differences between data were analysed in Prism (GraphPad Software) with appropriate post hoc multiple comparisons test. Tests are specified in figure legends.

## Acknowledgements

We thank Adam Hurlstone, Iain Hagan and all the members of the Cell Signalling lab for their critical reading of the manuscript, helpful comments and support. The Bioimaging Facility microscopes used in this study were purchased with grants from BBSRC, Wellcome Trust and the University of Manchester Strategic Fund. Special thanks go to Peter March for his help with microscopy. We thank the Molecular Biology Core Facilities and the Advanced Imaging Group, especially Steve Bagley and Kang Zeng at the Cancer Research UK Manchester Institute for their assistance with sequencing and microscopy.

## Author contributions

Conceptualization: A.P.P, E.L.O, H.J.W, A.M; Methodology: A.P.P, E.L.O, H.J.W; Validation: A.P.P., G.R.M.W.; Formal analysis: A.P.P, E.L.O; Investigation: A.P.P, G.R.M.W., E.L.O.; Resources: A.P.P, G.R.M.W., H.J.W; Writing - original draft preparation: A.P.P, A.M.; Visualization: A.P.P; Supervision: A.M.; Project administration: A.P.P, A.M.; Funding acquisition: A.P.P., A.M.

## Supplementary Materials

**Supplementary Figure 1.**
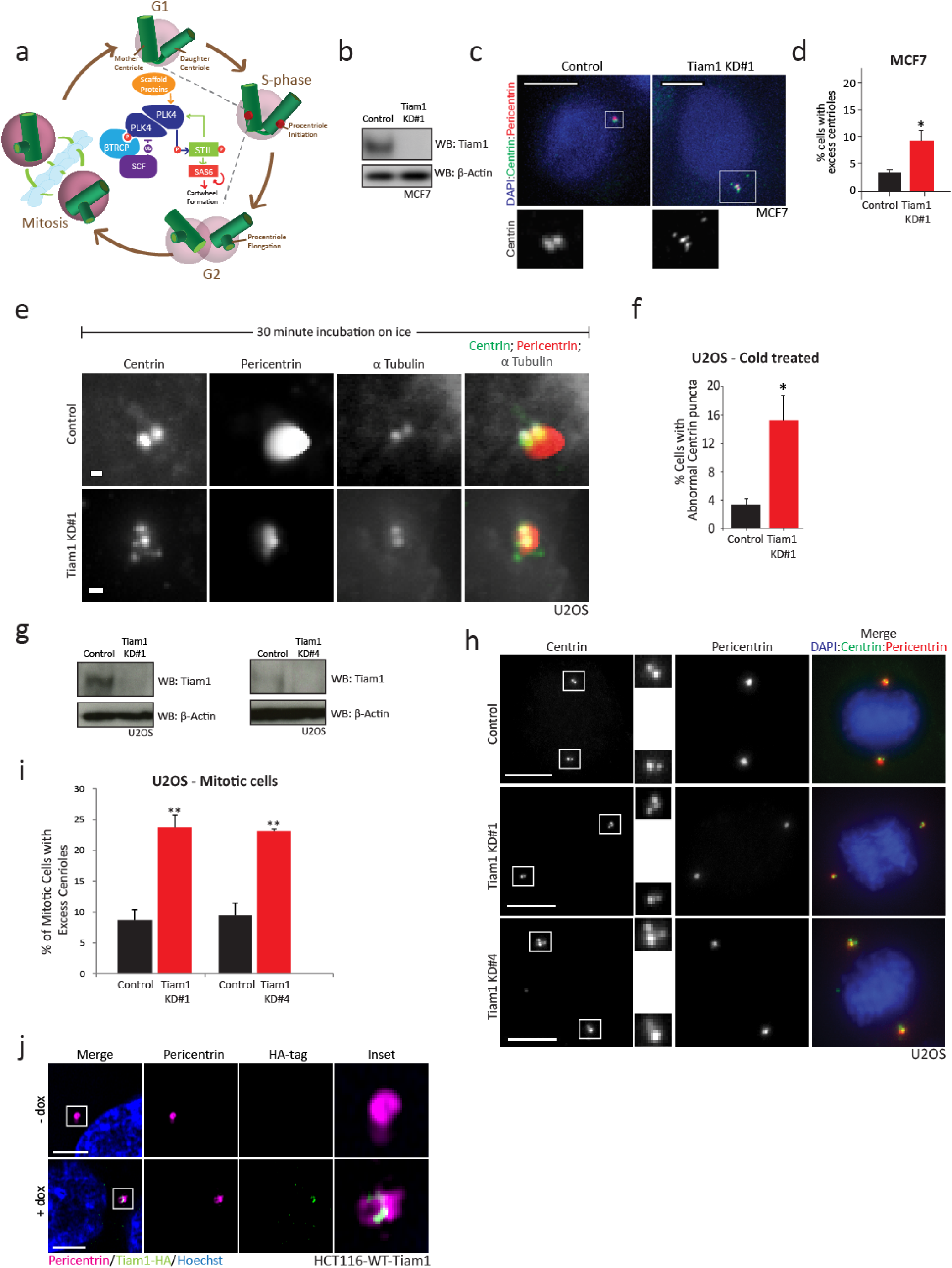
a) Schematic of the centriole duplication cycle. b) Western blot showing knockdown of endogenous Tiam1 in MCF7 cells. c) Deltavision maximal z-projection images of MCF7 cells showing an increase in centriole number following Tiam1 knockdown. Scale bar shows 10 μm. d) Quantification of centriole number in MCF7 cells as in (c), from three independent experiments, >145 cells quantified per experimental replicate. e) Deltavision images of centrosomes in U2OS cells (marked with Pericentrin), showing stable centriole staining (as marked with Centrin) in both control and Tiam1 knockdown cells, following cold treatment (see Methods). Scale bars show 1μm. f) Quantification of centriole number in control and Tiam1 knockdown cells following cold treatment as in (e), from three independent experiments. ≥70 cells were quantified per experimental replicate. g) Western blots showing knockdown of Tiam1 in U2OS cells following treatment with either Tiam1 KD#1 or Tiam1 KD#4 siRNAs. h) Deltavision maximal z-projection images of mitotic U2OS cells treated with either control or Tiam1 KD#1 or KD#4 siRNA, showing an increase in centriole number in Tiam1 knockdown cells. Scale bars show 10 μm. i) Quantification of cells with excess centrioles from images as in (h) from three independent experiments (Tiam1 KD#1; 100 mitotic cells quantified per experimental replicate; Tiam1 KD#4; ≥45 mitotic cells quantified per experimental replicate; N=3). * = p<0.05 ** = p<0.01. j) Single confocal z-planes of HCT116-Tiam1-WT cells showing centrosomal localisation of exogenous Tiam1 (as detected with an antibody against the HA tag on WT-Tiam1) following doxycycline (+dox) treatment. Scale bars show 3 μm. All bars show mean +/-SEM. β-Actin was used as a loading control in all blots.

**Supplementary Figure 2.**
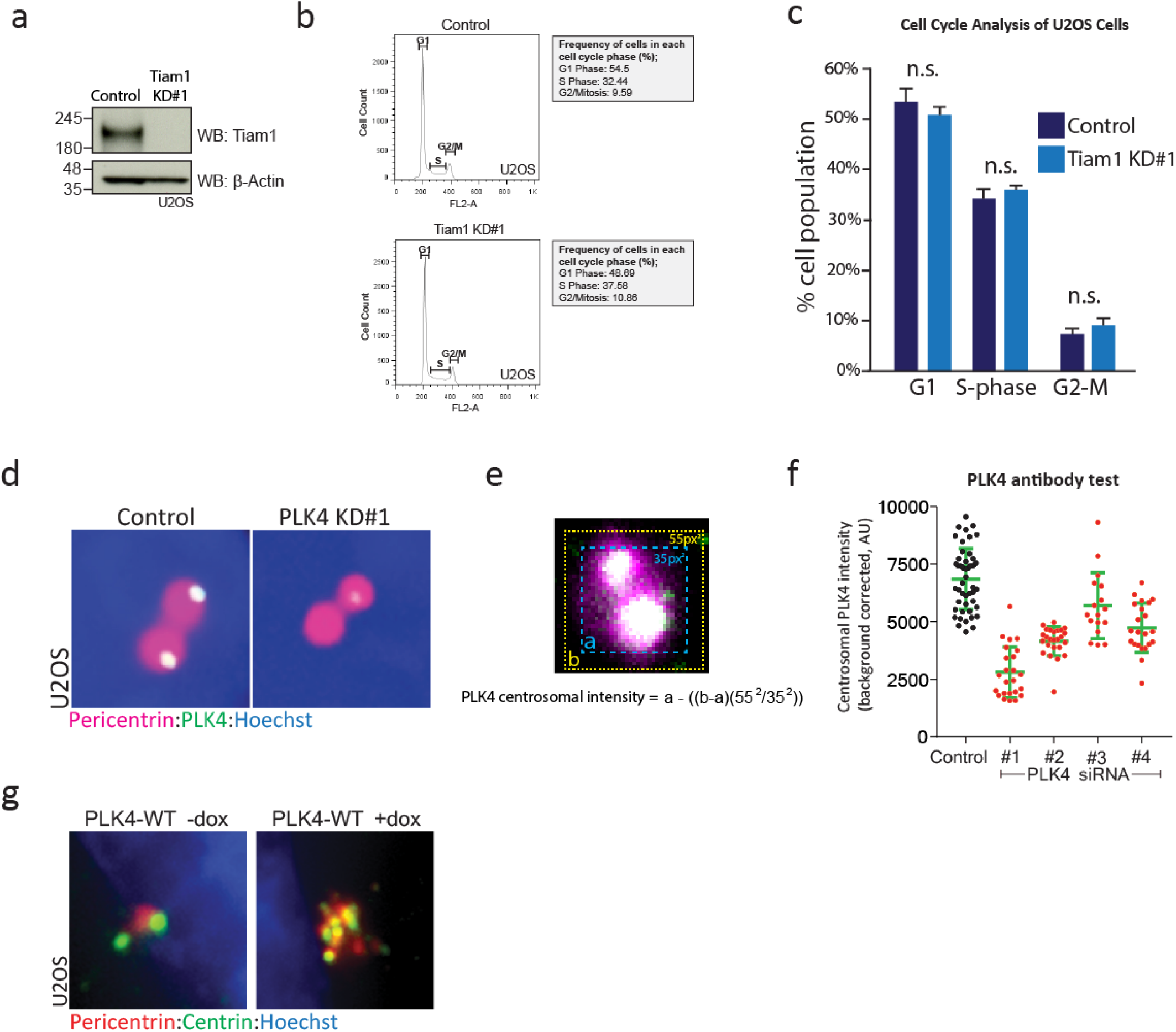
a) Western blot showing Tiam1 knockdown in U2OS cells. β-Actin was used as a loading control. b) Representative FACS plots from control and Tiam1 KD#1 U2OS cells. c) Summary of FACS analysis of cell cycle data from U2OS cells showing no change in cell cycle progression following Tiam1 knockdown, compared with control cells. Taken from three independent FACS experiments. d) Representative images of U2OS centrosomes (marked with Pericentrin), showing PLK4 staining at centrosomes in control cells, and decrease in staining in PLK4-knockdown cells. e) Schematic of centrosomal PLK4 intensity quantification. f) Quantification of centrosomal PLK4 intensity from U2OS cells treated with a panel of siRNAs targeting PLK4. g) Images of U2OS cells stained with Pericentrin (red, marking whole centrosomes) and Centrin (green, marking centrioles) showing an increase in centrosome and centriole number after treatment with doxycycline (+dox) to induce expression of WT-PLK4. t-tests comparing control and Tiam1 KD#1 for each cell cycle phase. n.s. = not significant

**Supplementary Figure 3.**
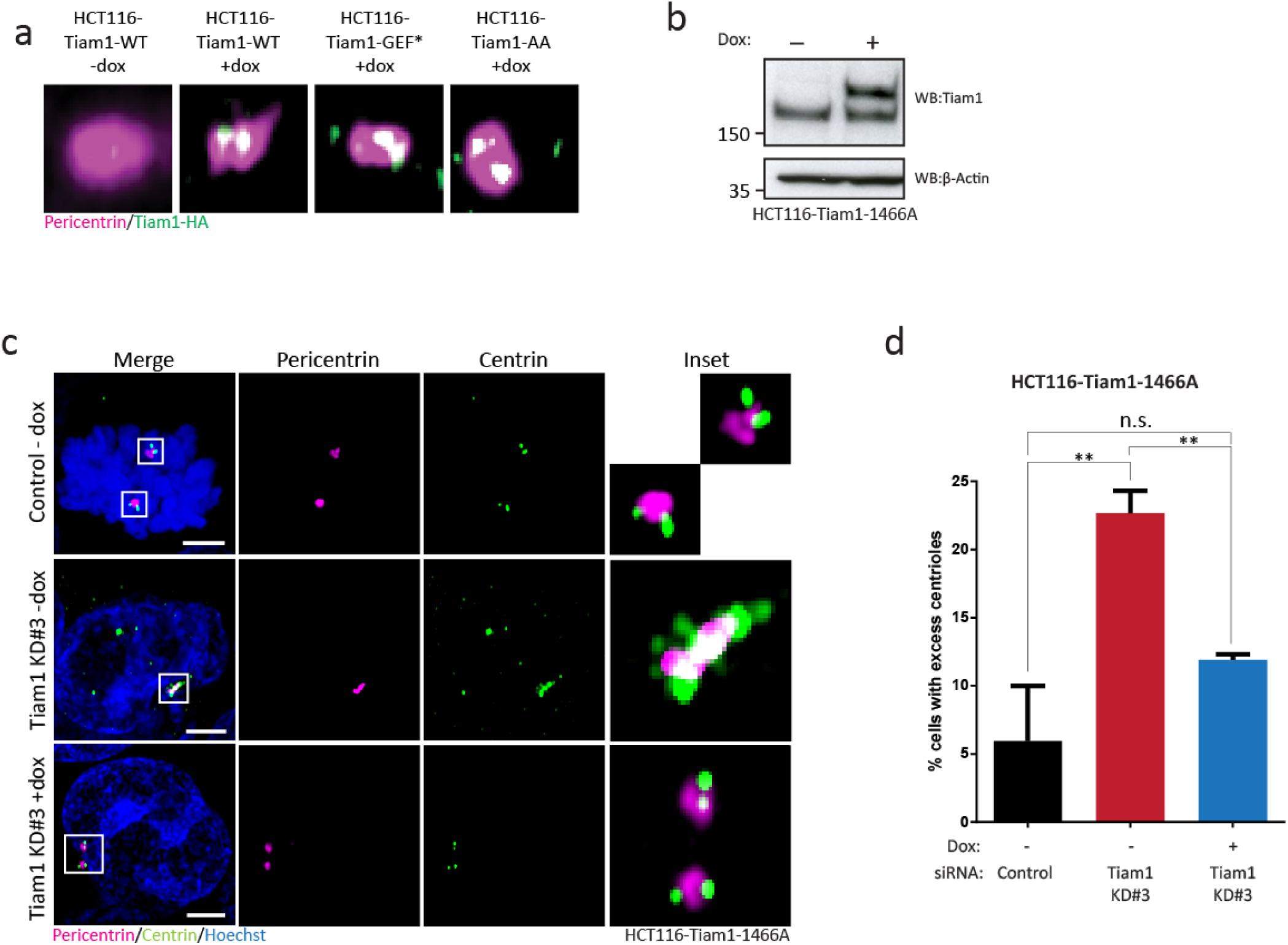
a) Confocal images of centrosomes (marked with Pericentrin, magenta) in HCT116 cells, showing localization of exogenous WT, GEF* and AA-mutant Tiam1 to the centrosome (marked with an antibody against the HA-tag, green.) b) Western blot showing expression of Tiam1-1466A in HCT116 following treatment with doxycycline (dox). β-Actin was used as a loading control. c) Maximal z-projection confocal images of HCT116 cells treated with either control or Tiam1 KD#3 siRNA, showing an increase in centriole number following Tiam1 knockdown, which is rescued by expression of Tiam1-1466A following doxycycline (Dox) treatment. d) Quantification of HCT116 cells with excess centrioles from cells as in (c) from 3 independent experiments; more than 50 cells counted per condition per experiment (one-way ANOVA, corrected for multiple comparisons). ** p<0.01 n.s. = not significant; error bars show S.E.M. Scale bars show 3μm.

**Supplementary Figure 4.**
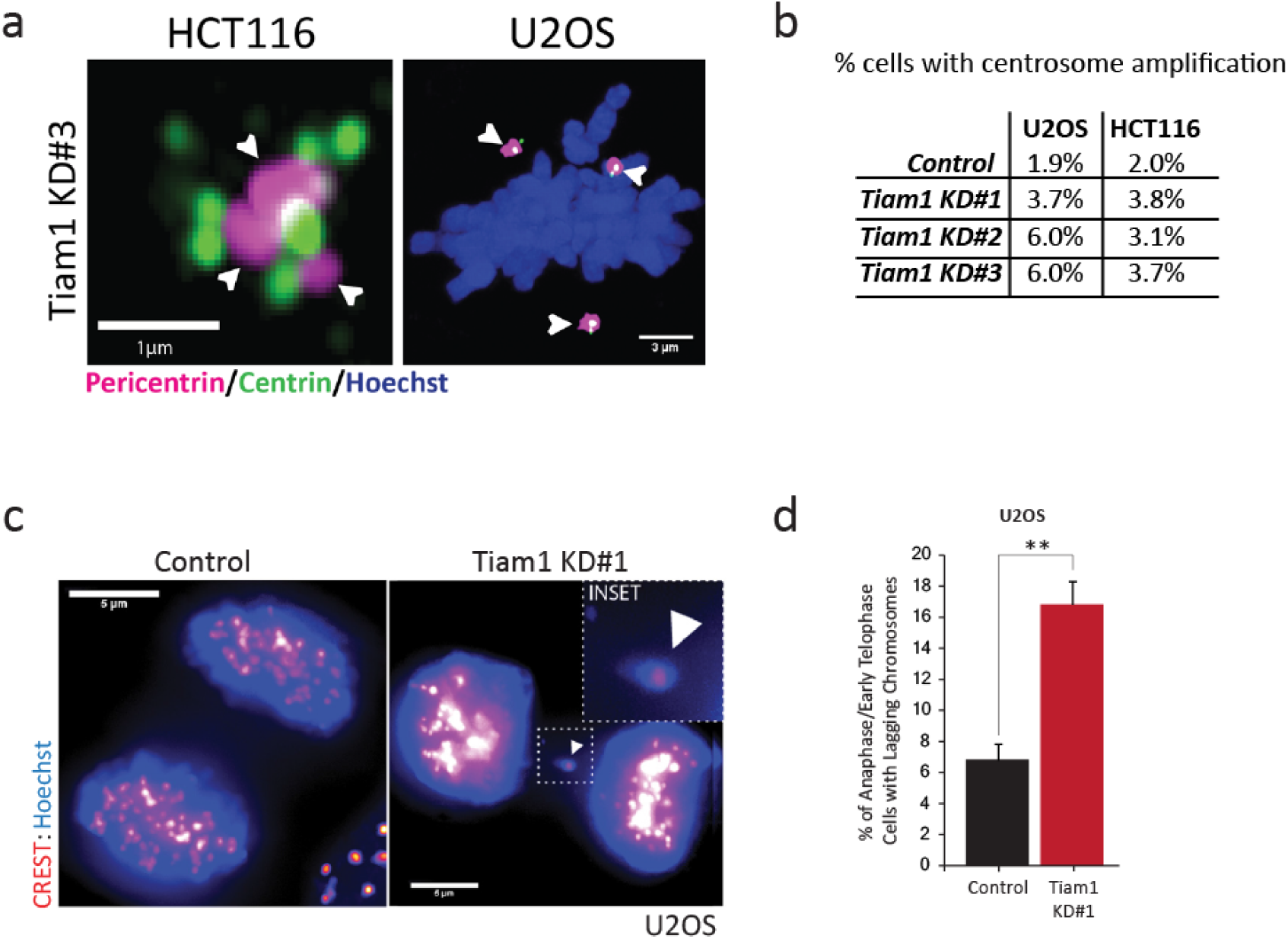
a) Representative confocal images showing HCT116 and U2OS cells with centrosome amplification (marked with arrowheads) following Tiam1 knockdown. Centrosomes are marked with Pericentrin (magenta) and Centrin (green). Scale bars are 1μm (HCT116) and 3μm (U2OS). b) Table showing increase in percentage of U2OS and HCT116 cells with excess centrosomes following three days of transfection with either control or Tiam1 knockdown siRNAs. c) Images of U2OS cells (stained with Hoechst and CREST as a centromere marker), with a lagging chromosome at telophase in the cell treated with Tiam1 KD#1 siRNA (marked with an arrowhead). Scale bars are 5μm. d) Quantification of control and Tiam1 knockdown U2OS cells with lagging chromosomes at anaphase and early telophase from three independent experiments. ** p<0.001 (t-test); error bars show S.E.M.

